# Fiber-parenchyma trade-off underlies changes in tropical forest structure and xylem architecture across a soil water gradient

**DOI:** 10.1101/2022.01.13.476164

**Authors:** Jehová Lourenço, Paulo Roberto de Lima Bittencourt, Brian J. Enquist, Camilla Rozindo Dias Milanez, Georg von Arx, Kiyomi Morino, Luciana Dias Thomaz, Lucy Rowland

## Abstract

- Wood anatomical traits can underpin tropical forest structural and functional changes across soil water gradients and therefore could improve our mechanistic understanding of how plants adapt to environmental change.
- We assessed how the variation in the forest maximum height (Hmax), stem diameter, and wood density (WD) is associated with variation in xylem traits (area of fibers and parenchyma, conductive area [CondA, sum of all vessels lumens], vessel lumen area [VLA], vessel density [VD], and vessel wall reinforcement [VWR]) across 42 plots of a Brazilian Atlantic Forest habitat that span strong soil water gradients.
- We found that in wetter communities, greater height and lower WD were associated with greater parenchyma area (capacitance), and lower fibers, VD, VWR. Contrastingly, in drier communities, lower height was associated with higher fiber area (xylem reinforcement), WD, VD, and VWR, while parenchyma area and vessels are reduced.
- Tree communities vary from conservative resource-use and structurally dependent hydraulic safety (Fibers) to acquisitive resource-use and capacitance dependent hydraulic safety (parenchyma). Such a fiber-parenchyma trade-off (FPT) underlies the variation in tree height across a soil water gradient. Wood anatomy is fundamental to understanding and predicting the impacts of environmental change on forest structure.

## Background

Size is one of the strongest sources of biological variation in forests, reflecting differences in plant form and function (Díaz *et al*., 2016), and adaptation to resource availability (e.g., soil water and nutrients) (Enquist, 2002; Lourenço Jr. *et al*., 2021). Understanding the intrinsic (anatomical) traits that underlie changes in plant size across soil water gradients may shed light on possible mechanisms critical for tree communities’ adaptation (Keddy, 1992) to environmental change. Ultimately, a more integrative understanding of how the effects of environmental change scale from xylem traits to individual trees and tree communities may improve our predictions of how tropical forests will respond to extended periods of increased temperature and droughts expected with climate change (IPCC, 2021).

Recent theoretical and empirical advances in plant allometry and anatomy have provided a more comprehensive and integrative understanding of the scaling relationship between plant size and the xylem architecture (West *et al*., 1999; Carrer *et al*., 2014; Rosell *et al*., 2017; Olson *et al*., 2020a). It has been demonstrated that tip-to-base widening of vessels is an essential anatomical adjustment to allow trees to grow taller by compensating for the increased hydraulic path length (Enquist, 2002; Olson *et al*., 2020a). Vessel widening exponentially increases the xylem water transport capacity at a rate of the fourth power of the conduit lumen diameter (Tyree & Zimmermann, 2002; Savage *et al*., 2010). Hence, larger conduits should be found in plants with more acquisitive strategies of resource use (e.g., high photosynthetic capacity) (Westoby, 1998; Wright *et al*., 2004) and low xylem construction costs (Eller *et al*., 2018) ultimately influencing forest structure along soil water gradients.

However, variation in tree size and hydraulic architecture requires anatomical changes beyond just vessel dimensions (Venturas *et al*., 2017; Lourenço Jr. *et al*., 2022) to facilitate plant adaptation to cope with environmental constraints (Chave *et al*., 2009). For example, environments poor in water and nutrients are known to select for low-statured plants, constraining the maximum tree height within plant communities (Hernández-Calderón *et al*., 2014; Scholten *et al*., 2017; Lourenço Jr. *et al*., 2021; Ohdo & Takahashi, 2021). Such variation in tree height may reflect trade-offs in traits underpinning resource use and hydraulic adaptations in plants (e.g., carbon allocation and water transport efficiency). A reduction in plant size may reflect investments in denser and more expensive tissues, increasing the organs’ life span (e.g., leaves and stems) where resource availability for tissue replacement is limited (e.g., well-drained and nutrient-poor soils) (Chave *et al*., 2009). It may also reflect the need for more drought-tolerant xylem hydraulic traits, which are often opposed to the traits needed to facilitate greater tree height, such as wider vessels (Olson *et al*., 2020b).

Xylem reinforcement via fibers and vessel walls increases wood density and has been linked to preventing vessel implosion under strongly negative xylem water potentials (Hacke *et al*., 2005; Ziemińska *et al*., 2013), as well as providing the mechanics to support the plant body against the bending forces (e.g., wind and gravity) (Niklas *et al*., 2006). Fiber area is hypothesized to trade-off with the area of parenchyma and mean vessel lumen area, due to mechanical constraints imposed by the wood space (Bittencourt *et al*., 2016), as predicted by the packing limits hypothesis (Baas, 1986; Schuldt *et al*., 2016). These trade-offs should have biomechanical (e.g., stem rigidity vs. flexibility) and physiological consequences (Niklas *et al*., 2006), reducing the amount of water transported (vessels) and stored (parenchyma) within the xylem, hence limiting photosynthesis and plant growth (Galmés *et al*., 2007).

Here, we highlight the hydraulic aspect of the fiber-parenchyma trade-off (FPT), which may reflect different hydraulic safety strategies in plants, via its effect on 1) fiber content influencing drought tolerance and; 2) parenchyma volume conferring greater hydraulic capacitance (Meinzer *et al*., 2009). High hydraulic capacitance increases hydraulic safety in the short term by buffering the fluctuations in xylem tension via the release of water from parenchyma to the vessels (Vesala *et al*., 2003). Ultimately, the FPT should result from different resource-use and hydraulic adaptations in plants, contrasting smaller plants with more conservative resource-use, higher fiber fraction conferring long-term hydraulic safety and biomechanical strength through greater wood density; and taller plants with more acquisitive resource-use, lower wood density, and shorter-term capacitance-dependent hydraulic safety (greater parenchyma area and vessels). Due to the link of the FPT to resource-use strategy, it is likely to predict community structure and height.

We assessed how the steep variation in forest structure (e.g., maximum tree height) across a gradient of soil water availability (Lourenço Jr. *et al*., 2021) is associated with changes in the xylem architecture in an environmentally diverse coastal habitat of the Brazilian Atlantic Forest (*restinga*). We hypothesize that 1) fiber and parenchyma area are the main drivers of xylem anatomical variability in topographical gradients of soil water availability; 2) the local variation in the forest structure (e.g., tree height) across soil water gradients will be associated with a trade-off between fibers and parenchyma, reflecting resource-use and hydraulic adaptation in plants. Hence, taller tree communities will have greater parenchyma area and larger vessels (hydraulic capacitance and efficiency), while smaller tree communities will have greater fiber area, denser wood, and thicker vessel walls (hydraulic safety via structure).

## Methods

### Study site

This study was conducted in a patch of restinga forest, a coastal habitat of the Atlantic Forest in the Espirito Santo State, Brazil (Fig. S1a). The study site is within the protected area of the Paulo Cesar Vinha state park (Fig S1b), where we set up 42 plots of 5×25m (Fig. S1c). The plots were positioned to capture fine spatial scale changes in water table depth – from floodable (valley regions with wetter soils), to intermediate (slight slope variation between valley and plateau) and drier regions (plateau regions where soils are sandier). More details on the forest inventory and the water table measurement can be found in Lourenço Jr. *et al*. (2021).

### Wood sampling and anatomical measurements

We collected wood samples from branches of at least 3 individual trees per species and per environment (floodable, intermediate, and drier sites). Thus, if a species occurred in a different environment, we collected their individuals again. For *Solanum sycocarpum* (n=1), *Buchenavia tetraphylla*, and *Campomanesia guazumifolia* (n=2), we collected a fewer number of individuals due to their rarity. The sampling was standardized to the first basal branch and approximately 2 meters from the tip to control for the tip-to-base scaling relationships (Olson *et al*., 2020a). We sampled adult trees, ranging from 5 to 20m height, and stem diameter>5cm. The anatomical measurements included only the 50% outermost region of the xylem area. The wood samples were sliced (10-20 μm), using a sledge microtome (Gärtner *et al*., 2014). Afterward, the samples were dyed in safranin (1%), and mounted in non-permanent slides (Gartner & Schweingruber, 2012). High-resolution images were taken in an Olympus BX 43 microscope using a 10× objective with an Olympus SC 30 camera coupled, which resulted in a resolution 248 of 1.546 pixels·μm-1. All images were analyzed with the ROXAS v3.0.560 (von Arx & Carrer, 2014) and the Image-Pro Plus v7.0 software (Media Cybernetics, Silver Spring, MD, USA). Many vessels were measured (170±113 cells) per image.

### Analytical framework

To assess the changes in community trait composition, we calculated the community weighted mean trait values (CWM) based on the species mean trait values weighted by the species abundance in each forest plot. Thus, the CWM for all analyzed traits (Table 1) was calculated for the 42 plots in our study area. All CWM trait values were used in the correlation analysis and for parameterizing the piece-wise structural equation modeling (SEM) to test for the possible relationships between the variation in forest maximum tree height (Hmax) and xylem anatomical traits. We used the package piecewiseSEM, version 1.2.1 (Lefcheck, 2016). We evaluated different possible effects of Hmax variation on xylem traits, and the best model was selected based on the lowest AICc and the highest P-value (Shipley, 2013). Only the selected model is shown. The analyses were done in R software (R Core Team, 2020). More information about the CWM and SEM analysis can be found in the supplementary material.

**Table 1.** Overview of wood anatomy traits, their acronym, units, and calculation methods description.

| Acronym | Unity | Trait | Description |
| --- | --- | --- | --- |
| Hmax | m | Maximum tree height | The mean value of the three tallest individuals per species in a community is calculated and then considered for the community weighted calculations |
| Stem<sub>d</sub> | cm | Stem diameter | The stem diameter at the breast height |
| WD | g cm<sup>-3</sup> | Wood density | Mass of wood per volume. |
| VD | no. mm<sup>-2</sup> | Vessel density | Number of vessels per area. |
| VLA | μm<sup>2</sup> | Vessel lumen area | Mean lumen area of vessels. |
| VWR | – | Vessel wall reinforcement | (t/b)<sup>2</sup>, where “t” is the double cell wall thickness and “b” the length of the same cell wall. |
| Fiber | % | Fiber Area | Proportion of the xylem cross-sectional area filled by fibers. |
| Parenchyma | % | Parenchyma area | Proportion of the xylem cross-sectional area filled by Parenchyma. |
| CondArea | % | Conductive area. Sum of vessel lumen areas | Proportion of the xylem cross-sectional area filled by the vessel lumen. |

## Results

### The fiber-parenchyma trade-off and forest structural variability across a soil water gradient

The increase of plant communities maximum height (Hmax) toward wet sites is associated with an increase in vessel lumen area (VLA) and parenchyma, and a reduction in fibers (Fig. 1a). The decrease in Hmax toward drier environments is associated with a greater investment in fibers (up to 80%), and a reduction in parenchyma (25 to 12.5%) and vessel lumen area (VLA, from 4000 to 1000µm^2^), while the conductive area (CondA, sum of all vessel lumens) is virtually invariable (from 10% to 7.5%) (Fig. 1a,b). Variation in fibers strongly trades-off with parenchyma (*R*^*2*^=0.93, *Slope*=0.90). These modifications in the proportion of xylem tissues and VLA are easily observed in the anatomical images (Fig. 1c).

**Figure 1.**
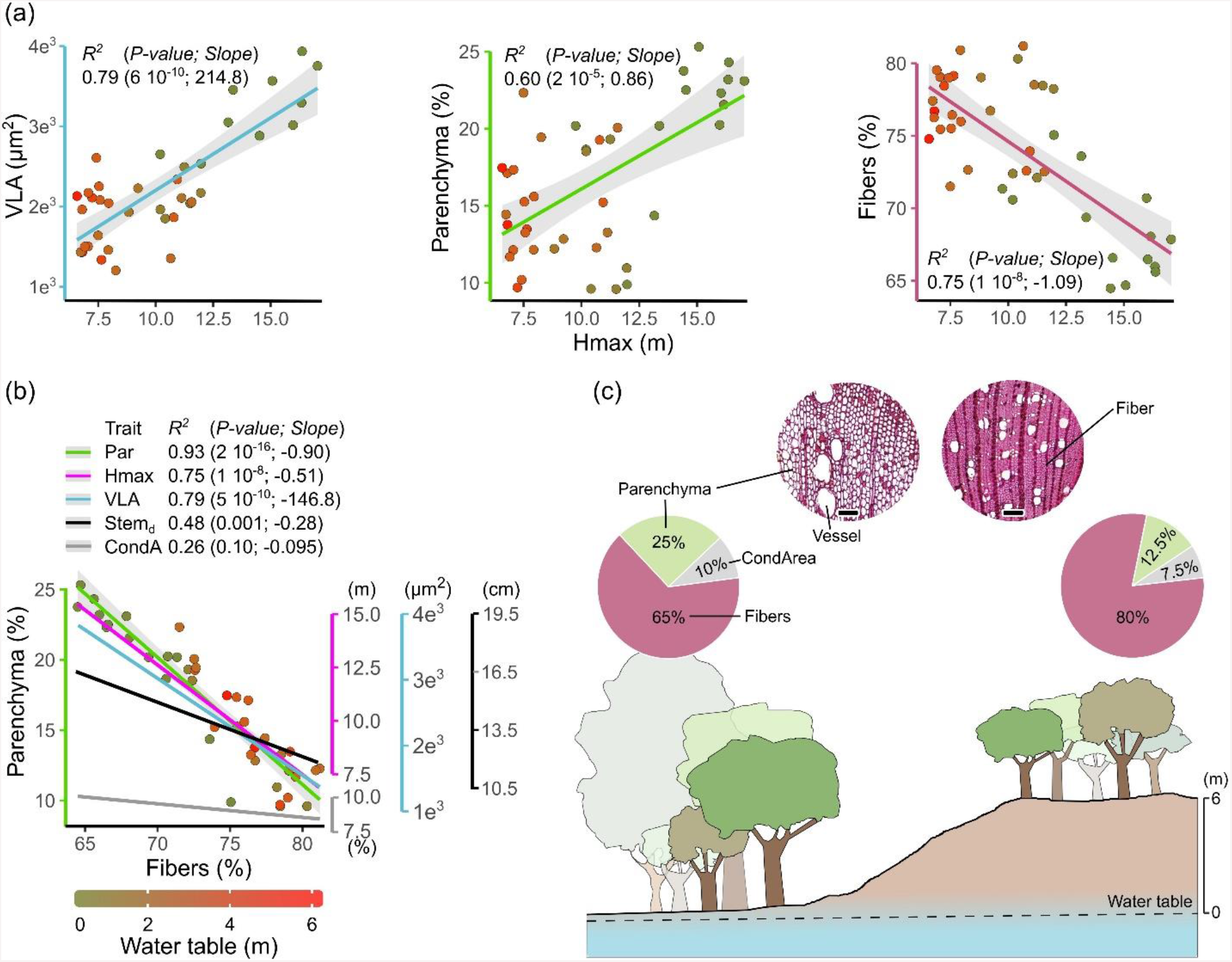
(a) Correlations between the variation in maximum tree height (Hmax) and the variation in vessel lumen area (VLA), parenchyma and fibers area. (b) The trade-off between fibers and parenchyma is mirrored by changes in Hmax, stem diameter (Stem_d_), VLA and conductive area (CondA, sum of all vessels’ lumen). Each dot represents a forest plot. (c) Scheme of the study site showing the variation in forest structure and xylem traits across the water table. Wood anatomy images are *Pseudobombax grandiflorum* (left) and *Pouteria coelomatica* (right). Scale bar represents 100 µm.

### The multiple effects of the fiber-parenchyma trade-off on xylem architecture and forest structure

The variation in Hmax caused by soil water gradient drives multiple changes in xylem traits and forest structural characteristics, via its direct effect on fibers (Fig. 2, and Table S1 – supplementary material). The per-unit reduction of parenchyma (−1.03) and VLA (−0.46) with increasing fibers fraction were the strongest relationships in the SEM, which directly explained the changes in height across the environmental gradient (Fig. 2a). Traits related to hydraulic safety and xylem structural reinforcement, such as vessel density (VD), wood density (WD), and vessel wall reinforcement (VWR), were negatively associated with tree height, most likely due to their positive link to fiber area toward drier environments (Fig. 2a). In contrast, traits associated with hydraulic capacitance and hydraulic efficiency (parenchyma area and VLA), were found in wetter and taller tree communities.

**Figure 2.**
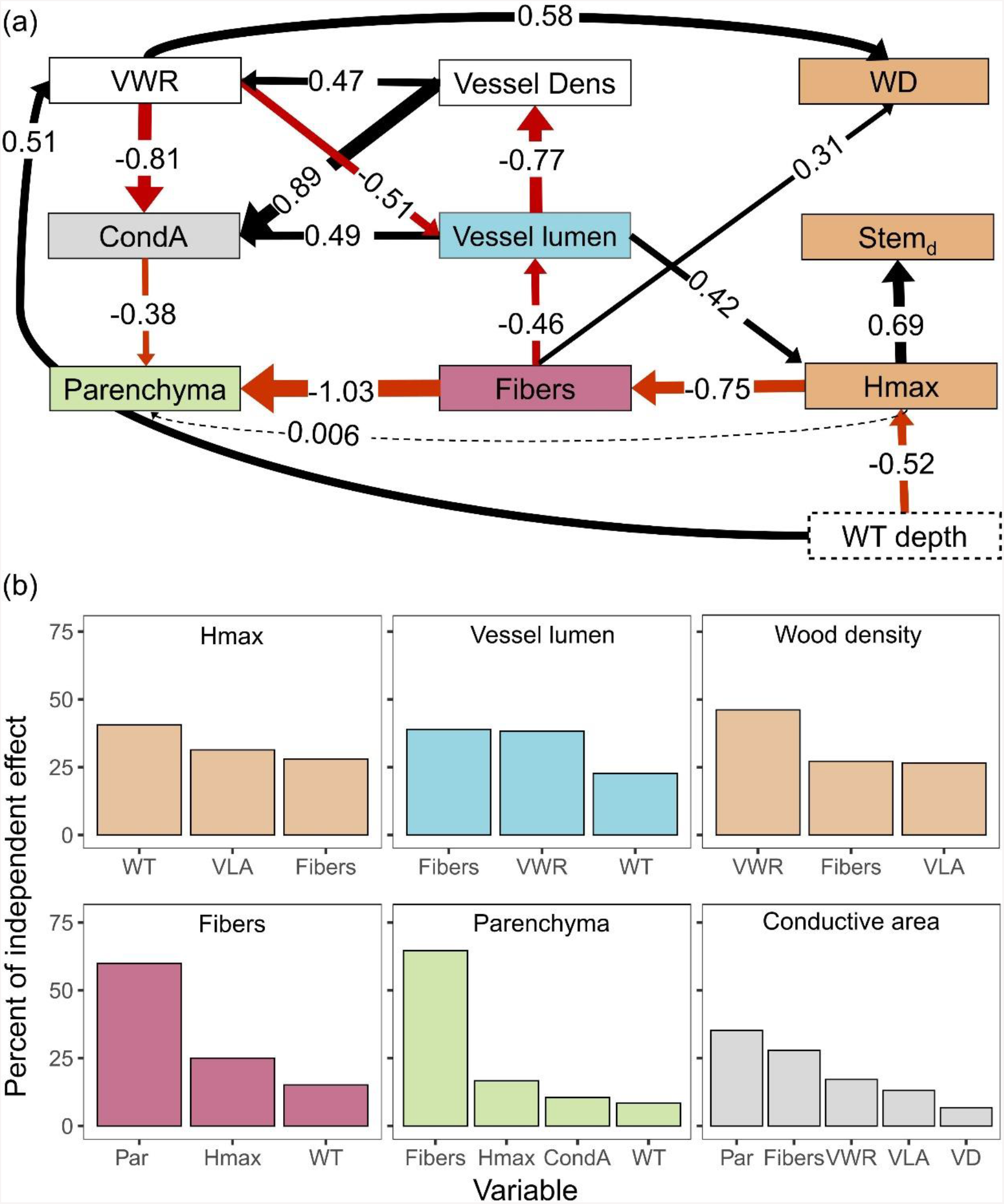
(a) Piecewise structural equation modeling (SEM) analysis represented by path diagrams exploring the xylem traits underlying the variation in maximum tree height (Hmax), stem diameter (Stem_d_), and wood density (WD) across a soil water table gradient (WT). (b) The percent of the independent effects of different predictors are shown. VWR - vessel wall reinforcement, Vessel Dens – vessel density, CondA – conductive area, Par – parenchyma, and VLA – vessel lumen area.

## Discussion

### Variation in tree height across a soil water gradient is associated with a trade-off between fibers and parenchyma, reflecting differences in resource use and hydraulic adaptation

The negative association between Hmax and fiber fraction reveals that plants in drier regions adopt a more conservative strategy of resource use and growth, investing in more perennial (denser) wood tissues (Chave *et al*., 2009). High investment in fibers and vessel walls should confer hydraulic safety in these drier tree communities (Hacke *et al*., 2001; Meinzer *et al*., 2009), reinforcing the xylem against the likely more negative water potentials. This reinforcement of the xylem should cause mechanical constraints in the wood space, restricting both vessel size and parenchyma area (Fig. 1b), as predicted by the packing limit hypothesis (Baas, 1986; Schuldt *et al*., 2016). Hence, the constraining effects of fibers on vessels and parenchyma should result in physiological implications, for example, limiting the capacity of the wood to transport and store water, respectively. Meinzer *et al*. (2009) proposed that hydraulic safety in hardwood drought-adapted plants should primarily rely on the reinforcement of the xylem structure against negative water potentials (e.g., fibers), while short-term hydraulic safety in softwood plants adapted to wetter conditions should rely on hydraulic capacitance (e.g., parenchyma), which should buffer the transpiration-induced daily fluctuation of the water potential within the hydraulic system.

Thus, the variation in fibers and parenchyma (FPT) is likely to underlie the changes in forest structure. Tall trees should rely on greater hydraulic capacitance and conductivity (parenchyma area and vessel lumen), while small trees should rely on greater hydraulic safety and denser tissues (larger fibers area and denser wood). Ultimately, the FPT may reflect different strategies of resource use, growth, and hydraulic adaptation in plants, suggesting that variation in tree height may not be just dependent on vessel widening (Olson *et al*., 2020a), but more complex variations in the xylem architecture, coordinated by xylem trade-offs at the tissue level. The interaction of height, wood anatomy, and water availability should be a key area for future research.

### Fiber-parenchyma trade-off drives multiple changes in hydraulic architecture via a packing limit mechanism

Our results provide support for the hypothesis that the packing limits, the physical constraint in the wood space (Baas, 1986; Sperry *et al*., 2008; Schuldt *et al*., 2016), are driven by the trade-off between fibers and parenchyma. The variation in fibers has the strongest independent effects on parenchyma and VLA, underlying a chain of effects in other xylem traits that shape the hydraulic architecture of trees across the soil water gradient. The limitation of space within the wood due to the increase of fibers toward drier plant communities has direct and negative effects on VLA, and subsequent positive indirect effects on the vessel density and wall thickness. These modifications should strongly determine the ability of plants to adapt to limiting soil water conditions, as it is likely to reduce the amount of water transported (Tyree & Zimmermann, 2002; Hacke *et al*., 2006; Zhu *et al*., 2017; Liu *et al*., 2020), reinforce the xylem against strongly negative water pressure (Hacke *et al*., 2001) and increase the capacity of the plant to bypass embolized vessels (von Arx *et al*., 2013; Schuldt *et al*., 2016). These traits all favor more conservative strategies of resource use and growth (Wright *et al*., 2004; Chave *et al*., 2009; Díaz *et al*., 2016). In contrast, plant communities in wetter conditions experiencing less negative water potentials should require less investment in fibers and vessel walls (Bittencourt *et al*., 2016), allowing more wood space for vessels and parenchyma. These modifications will increase the hydraulic efficiency, favoring plants with more acquisitive and fast-growth strategies. The multiple changes in xylem traits driven by a fiber-parenchyma trade-off (FPT) reinforce the hypothesis that hydraulic adaptation in plants should rely on the variation of multiple traits (Venturas *et al*., 2017), underlying the local variation in forest structure.

## Acknowledgments

This work was partially financed by CAPES/Brazil (88881.131961/2016-01). The British Royal Society (Newton International Fellowship-NIF\R1\191168) currently supports this research. We thank the support of Dr. Malcolm Hughes, Instituto Estadual de Meio Ambiente (IEMA/ES), Parque Paulo César Vinha (PEPCV) staffs, Oberdan Pereira, Douglas Wandekoken, Felipe Barreto, Fabiano Volponi, Jocimara Lourenço, Nilton Filho, Rodrigo Theófilo, José Gomes, and Danielly Hirata.

## Author’s contributions

JLJ and CRD designed the project; LDT gave support to the taxonomy survey, wood collection, and plant species identification; KM gave support to the wood anatomy analysis. JLJ and GVA analyzed the data; JLJ, LR, PRLB, BJE, and GVA led the writing of the manuscript. All authors contributed critically to the drafts and gave final approval for the publication.

## Data accessibility

Additional Supporting Information may be found in the online version of this article:

## Supporting information

Doc file containing the map of the study site, additional analysis, and additional details of the statistical approach.

## Coding

Text file script to the community-weighted metrics and SEM analysis.

