## Supplementary material for "Fiber-parenchyma trade-off underlies changes in tropical forest structure and xylem architecture across a soil water gradient"

### Contents

- **Table S1.** Direct, indirect, and total effects between the main variables used in the structural equation modeling.
- **Figure S1.** Study site maps and experimental design.
- **Figure S2.** Correlation plots comparing variation in fibers with variation in xylem and forest characteristics
- **Community weighted calculations**
- **Variable selection criteria for SEM constructs**

**Table S1.** Piecewise structural equation models exploring the effects of Fibers, maximum tree height (Hmax), and water table (WT) on xylem traits and plant community structure.

| Predictor | Response variable | | | | | | | | |
|---|---|---|---|---|---|---|---|---|---|
| | a) Xylem traits | | | | | | b) Forest structural traits | | |
| | Fibers | Par | CondA | VLA | VD | VWR | WD | Hmax | $Stem_d$ |
| Fibers | **...** | **-1.02**, -1.03, 0.01 | **0.16**, NA, 0.16 | **-0.46**, -0.46, NA | **0.35**, NA, 0.35 | **0.16**, NA, 0.16 | **0.41**, 0.31, 0.10 | **-0.19**, NA, -0.19 | **-0.13**, NA, -0.13 |
| Hmax | **-0.75**, -0.75, NA | **0.77**, NA, 0.77 | **0.04**, NA, 0.04 | **0.34**, NA, 0.34 | **0.26**, NA, 0.26 | **0.12**, NA, 0.12 | **-0.30**, NA, -0.30 | ... | **0.69**, 0.69, NA |
| WT | **0.39**, 0.39, NA | **-0.40**, NA, -0,40 | **-0.37**, NA, -0.37 | **-0.40**, NA, -0.40 | **0.14**, NA, 0.14 | **0.57**, 0.51, 0.06 | **0.35**, NA, 0.35 | **-0.69**, -0.52, -0.17 | **-0.36**, NA, -0.36 |

Notes: Values are expressed as standardized effect sizes: total effect (in bold) followed by direct effect and indirect effect values. These parameters were directly assessed via the community-weighted means fibers, parenchyma (Par), conductive area (CondA, sum of all vessel lumens), mean vessel lumen area (VLA), vessel density (VD), vessel wall reinforcement (VWR) wood density (WD), tree maximum height (Hmax) and stem diameter ($Stem_d$) for 42 woody plant communities of restinga habitats of Atlantic forest. AICc is Akaike's information criterion estimated from Shipley's *d* separation test (Shipley 2013). *C*, df, *P*, and AICc values for variables: 40.99, 56, 0.93, and 530.99. An ellipsis, or "NA," indicates no effect between predictor and response variable.

**Figure S1.** (a) Map of Brazil highlighting the Espírito Santo State (ES), where (b) Paulo Cesar Vinha state park (PCV) and the study area are located (circle). (c) We set up 42 plots (measuring 5 x 25m) that span a steep soil water gradient caused by the microtopographic-driven variability in the water table in a short spatial scale of 207 ± 60 meters from floodable (○) to intermediate (□) and drier (Δ) forest communities. The water table is not related to seawater. The rainfall is the source of water into this system, causing periodic waterlogging in lower areas (valleys) more frequently in the rainy summer season. Such areas in *restinga* habitats are named floodable forests.

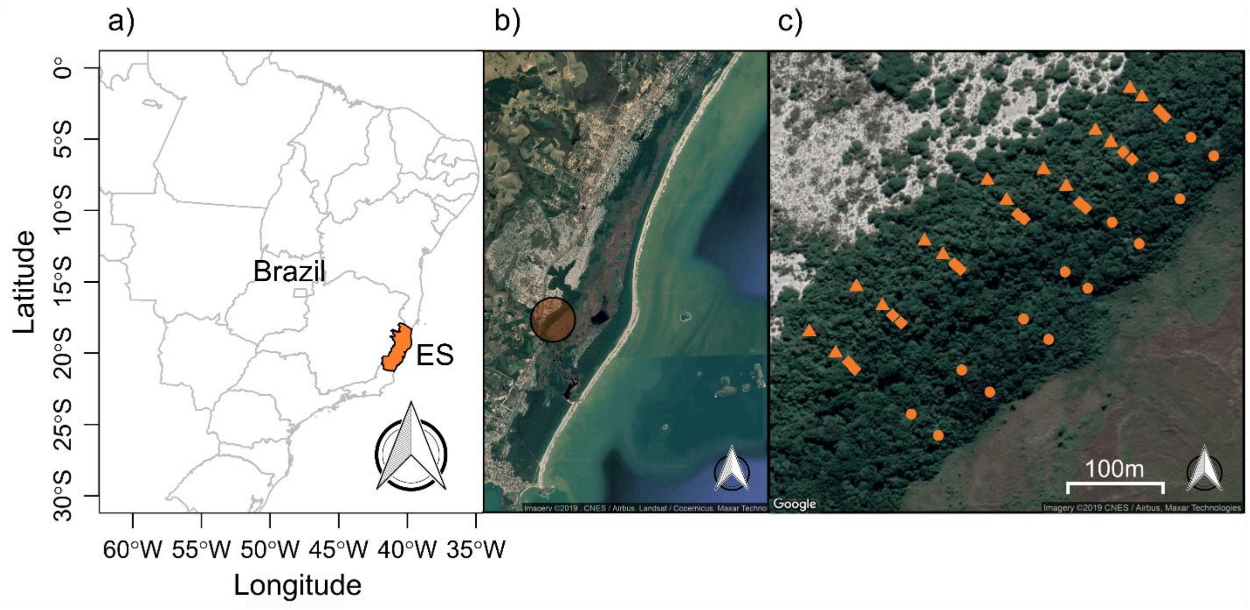

**Figure S2.** Correlation plots comparing variation in fibers with variation in xylem and forest characteristics.

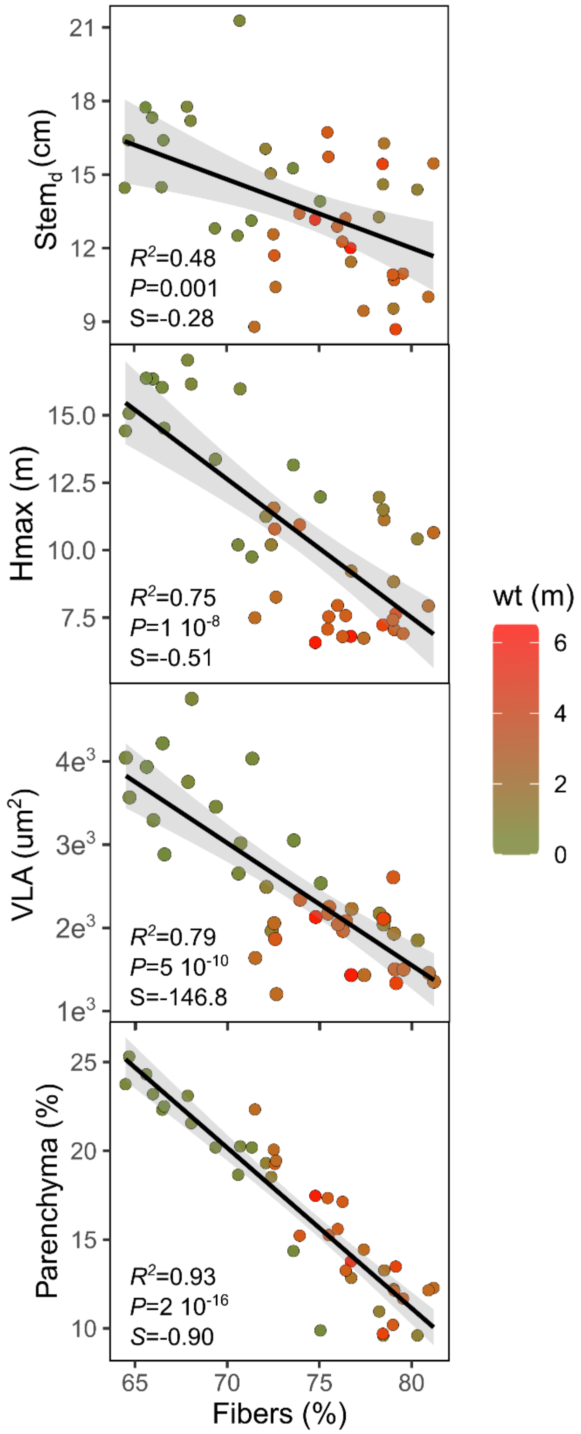

**Community weighted calculations**

We calculated the community-weighted mean (CWM) for each of the 42 plots (or plant communities, for simplicity) in this study. The calculations were made by the species trait values weighted by the species abundance, according to the equation:

$$CWM_{j,y} = \sum_{k=1}^{n_j} A_{kj}.z_k$$

where $n_j$ is the number of species sampled in plot j, $A_{k,j}$ is the relative abundance of species k in plot j, and $z_k$ is the mean value of species k.

**Variable selection criteria for SEM constructs**

As general criteria, the selection of the variables used in this study was based on their relevance in describing parameters of soil water availability (water table depth), forest structure (maximum tree height and stem diameter), and wood architecture (fibers, parenchyma, conductive area (CondA), vessel lumen area (VLA), vessel density (VD), vessel wall reinforcement (VWR), and wood density(WD)). The association between variables was based on the literature, as described in the introduction of the main text article, including theoretical and empirically demonstrated links between the variables. The variables associations are represented in the path analysis (Fig. 2a) and based on the structural equation modeling (SEM).
